# The Sigma-2 Receptor/TMEM97, PGRMC1, and LDL Receptor complex are responsible for the cellular uptake of Aβ42 and its protein aggregates

**DOI:** 10.1101/2020.02.19.956359

**Authors:** Aladdin Riad, Zsofia Lengyel-Zhand, Chenbo Zeng, Chi-Chang Weng, Virginia M.-Y. Lee, John Q. Trojanowski, Robert H. Mach

## Abstract

**Background:** Our lab has recently shown that the Sigma-2 Receptor/Transmembrane Protein 97 (TMEM97) and Progesterone Receptor Membrane Component 1 (PGRMC1) form a complex with the Low Density Lipoprotein Receptor (LDLR), and this intact complex is required for efficient uptake of lipoproteins such as LDL and apolipoprotein E (apoE). These receptors are expressed in the nervous system where they have implications in neurodegenerative diseases such as Alzheimer’s Disease (AD), where apoE is involved in neuronal uptake and accumulation of Aβ42, eventually cascading into neurodegeneration, synaptic dysfunction, and ultimately, dementia.

**Hypothesis:** We hypothesize that the intact Sigma-2 receptor complex –TMEM97, PGRMC1, and LDLR— is necessary for internalization of apoE and Aβ42 monomers (mAβ42) and oligomers (oAβ42), and the disruption of the receptor complex inhibits uptake.

**Results:** The results of this study suggest that the intact Sigma-2 receptor complex is a binding site for mAβ42 and oAβ42, in the presence or absence of apoE2, apoE3, and apoE4. The loss or pharmacological inhibition of one or both of these proteins results in the disruption of the complex leading to decreased uptake of mAβ42 and oAβ42 and apoE in primary neurons.

**Conclusion:** The TMEM97, PGRMC1, and LDLR complex is a pathway for the cellular uptake of Aβ42 via apoE dependent and independent mechanisms. This study suggests that the complex may potentially be a novel pharmacological target to decrease neuronal Aβ42 internalization and accumulation, which may represent a new strategy for inhibiting the rate of neurotoxicity, neurodegeneration, and progression of AD.

## INTRODUCTION

The sigma receptors are a family of receptors who are poorly characterized. They were initially thought to be a subset of the opioid receptor family, but were later identified to be a distinct subset of receptors that include the Sigma-1 and Sigma-2 receptors [1–4]. The Sigma-2 receptor is expressed highly in proliferating cancer tumor cells and has been an attractive target for imaging solid tumors [2]. The Sigma-2 receptor has also been used as a target for the treatment of central nervous system (CNS) disorders such as depression and psychosis [5], and it has been recently studied as a potential pharmacological target for treating Alzheimer’s Disease (AD) [6, 7].

Our previous work has shown that Sigma-2 Receptor/Transmembrane Protein 97 (TMEM97) and Progesterone Receptor Membrane Component 1 (PGRMC1) form a complex with the Low Density Lipoprotein Receptor (LDLR) [8], and this intact complex is important for the efficient uptake of low density lipoprotein (LDL). We found that disrupting this complex by genetically ablating or by pharmacologically targeting TMEM97 or PGRMC1 resulted in a decreased capacity for LDL internalization. For this reason, we choose to study the role of this complex in the internalization of Amyloid Beta 1-42 (Aβ42) and apolipoprotein E (apoE).

Aβ42 is a peptide formed from the cleavage of amyloid precursor protein (APP) [9–11]. The accumulation of Aβ42 in the brain results in aggregation into neurotoxic oligomers and fibrils [12] leading to deposition into amyloid plaques, the hallmark of a brain afflicted by AD [13]. Although amyloid beta plaques are extracellular, mounting evidence suggests that aggregation also begins within neurons, where uptake and subsequent degradation within lysosomes has been dysregulated leading to intraneuronal accumulation leading to aggregation [14–20]. Recent evidence suggests the cell-to-cell transfer of Aβ42 also plays a role in the pathogenesis of AD [20–25]. Reducing neuronal uptake may slow the cytotoxic effects of intracellular aggregation.

ApoE is a protein that mediates lipid homeostasis by facilitating transport of lipids within the various aqueous biological environment to cells and tissues within the body [26]. Within the CNS, apoE is primarily produced by astrocytes and transports cholesterol to neurons via its recognition by the LDLR family including LDLR and LDLR-related protein (LRP) receptors [27]. The main isoforms of apoE is apoE3, with apoE2 and apoE4 being the result of a single amino acid mutation. The presence of one apoE4 allele results in a 3-fold increased risk for developing AD, while the presence of two alleles result in a 12-fold increased risk factor, while patients with one or more apoE2 alleles have a reduced risk for developing AD [28]. ApoE has been shown to interact with Aβ42 and affect its clearance from the CNS, and increasing deposition into plaques [29].

In this study, we examined the role of the TMEM97/PGRMC1/LDLR protein complex in the uptake of Aβ42 and its protein aggregates and apoE, separately and when in a complex with one another. Our studies assessed the role of uptake in a HeLa cell model system, and in a primary neuronal cell culture system. We provide experimental evidence for the importance of this intact complex for the uptake of Aβ42 and apoE, and show that disrupting the complex pharmacologically results in decreased uptake, which alludes to the prospect that this complex being a therapeutic target for limiting the progression of AD.

## RESULTS

### The role of the TMEM97 and PGRMC1 in Aβ42 and apoE uptake

Samples of Aβ42 monomers (mAβ42), oligomers, (oAβ42), and fibrils were prepared using previously published methods [30–32]. Transmission electron microscopy (TEM) images showed that mAβ42 preparations contained no aggregated particles, oAβ42 samples showed small, round particles with no protofibrils or fibrils present, and fAβ42 samples contained well defined filaments with no protofibrils or aggregated oligomers (SI Appendix, Fig. S1A). Dot blot analysis indicated positive reactivity for all samples with Aβ42 6E10 antibody, which recognizes all species of Aβ42, however only oAβ42 showed positive reactivity to the oligomer specific A11 antibody, indicating pure samples of the three aggregation states (SI Appendix, Fig. S1B).

Our previous work indicated that TMEM97 and PGRMC1 form a trimeric complex with the LDLR, and this complex is necessary for efficient uptake of LDL. As apoE can be taken up through the LDLR and is associated with Aβ42, we sought to assess this complex’s role in uptake of Aβ42 and apoE3. To conduct these experiments we utilized four HeLa cell lines, control (Scramble/Cas9), TMEM97 KO, PGRMC1 KO, and TMEM97/PGRMC1 double KO (DKO). These cell lines were generated in our group utilizing CRISPR/Cas9 technology; previous studies characterized these cell lines and western blot data demonstrated a complete ablation of TMEM97 and/or PGRMC1 in the knockout cells [8, 33]. Cells were incubated in lipoprotein-depleted serum (LPDS) for 24 hours in order to induce expression of LDLR. Cells were treated with 1000 nM mAβ42, oAβ42, or fAβ42 alone and in combination with 250 nM apoE for 24 hours at 37 °C. Cell associated Aβ42 and apoE3 was quantified via ELISA (Fig. 1 A and B). The first observation of these experiments was mAβ42 is more efficiently taken up by cells when it is in a complex with apoE3. Uptake of the mAβ42/apoE3 complex was significantly decreased in the TMEM97 KO, PGRMC1 KO, and DKO cell lines when compared to Scramble/Cas9. Uptake of oAβ42, and to a lesser extent fAβ42, alone and when in complex with apoE3 was also significantly decreased in the knockout cell lines. Uptake of apoE3 was also significantly less in the knockout cell lines compared to Scramble/Cas9 control cell (Fig. 1B). These results indicate that TMEM97 and PGRMC1 mediate internalization of Aβ42 and apoE3, so our next goal was to evaluate the effect of pharmacological inhibition of TMEM97 and PGRMC1 on Aβ42 and apoE3 internalization in the Scramble/Cas9 cells. We utilized two TMEM97 ligands, RHM-4 (Kd ~ 0.2 nM) and SW43 (Kd ~ 12nM), and a PGRMC1 ligand AG-205 (Kd ~ 1 μM against Sigma2R [8]). Scramble/Cas9 cells when treated with 500 nM RHM-4, SW43, or AG-205 showed significantly reduced capacity for internalizing mAβ42 and oAβ42 alone or when complexed with apoE3; and to a lesser extent reduced the cell associated fAβ42 alone or when complexed with apoE3 (Fig. 1C). The compounds also resulted in significantly less uptake of apoE3 alone or when in a complex with any of the Aβ42 aggregated states (Fig 1D).

**Figure 1.**
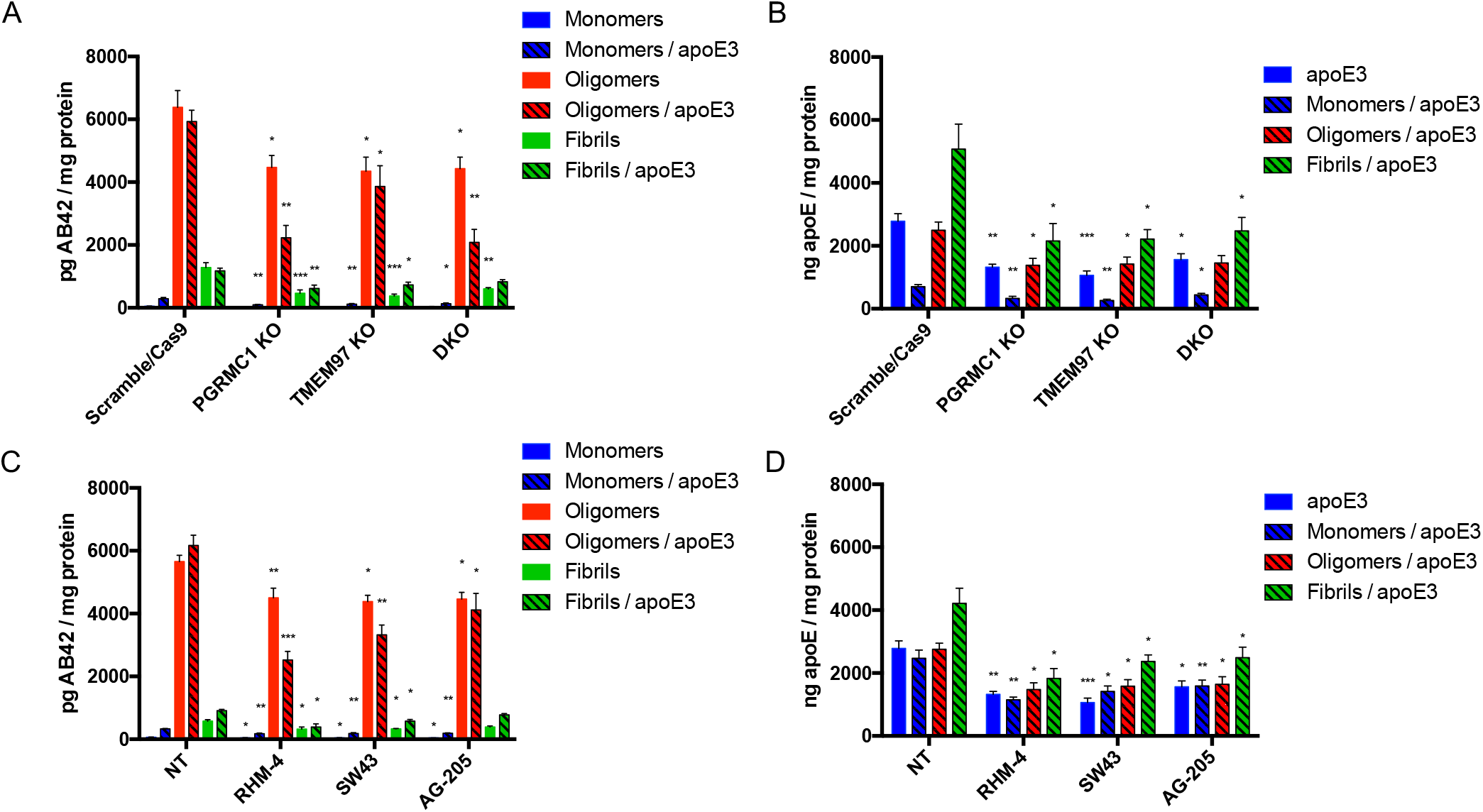
Disruption of the TMEM97/PGRMC1/LDLR complex reduces cell associated Aβ42 and apoE levels in HeLa cells. Cells were treated with 1000 nM Aβ42 monomers, oligomers, or fibrils in the presence or absence of 250 nM apoE for 24 hours at 37 °C. (A-B) Knockout cell lines compared to control cells (Scramble/Cas9) and cell associated (A) Aβ42 and (B) apoE was quantified. (C-D) Scramble/Cas9 cells treated with either 500nM RHM-4, SW43, or AG-205 were compared to no drug treated Scramble/Cas9 cells (NT) and cell associated (C) Aβ42 and (D) apoE was quantified. Cell associated Aβ42 and apoE was determined via ELISA of cell lysates normalized to total cell protein. Data represents mean ± SEM (n=3); *, p < 0.05; **, p < 0.01; ***, p < 0.001; **** p < 0.0001; one way ANOVA for each treatment compared to control.

To visualize uptake of oAβ42, the control and knockout cell lines were treated with 3 μM fluorescently labeled oAβ42 for 10 minutes, 30 minutes, and 60 minutes. Cells were washed thoroughly then imaged immediately after addition of Hoechst to stain nuclei. Scramble/Cas9 cells showed significantly more uptake of oligomers compared to the knockout cell lines at all time points since the internalized signal increased over time (Fig 2A). Interestingly fluorescent fAβ42 was not internalized by the cells, but accumulated on the cell surface (Fig 2B).

**Figure 2.**
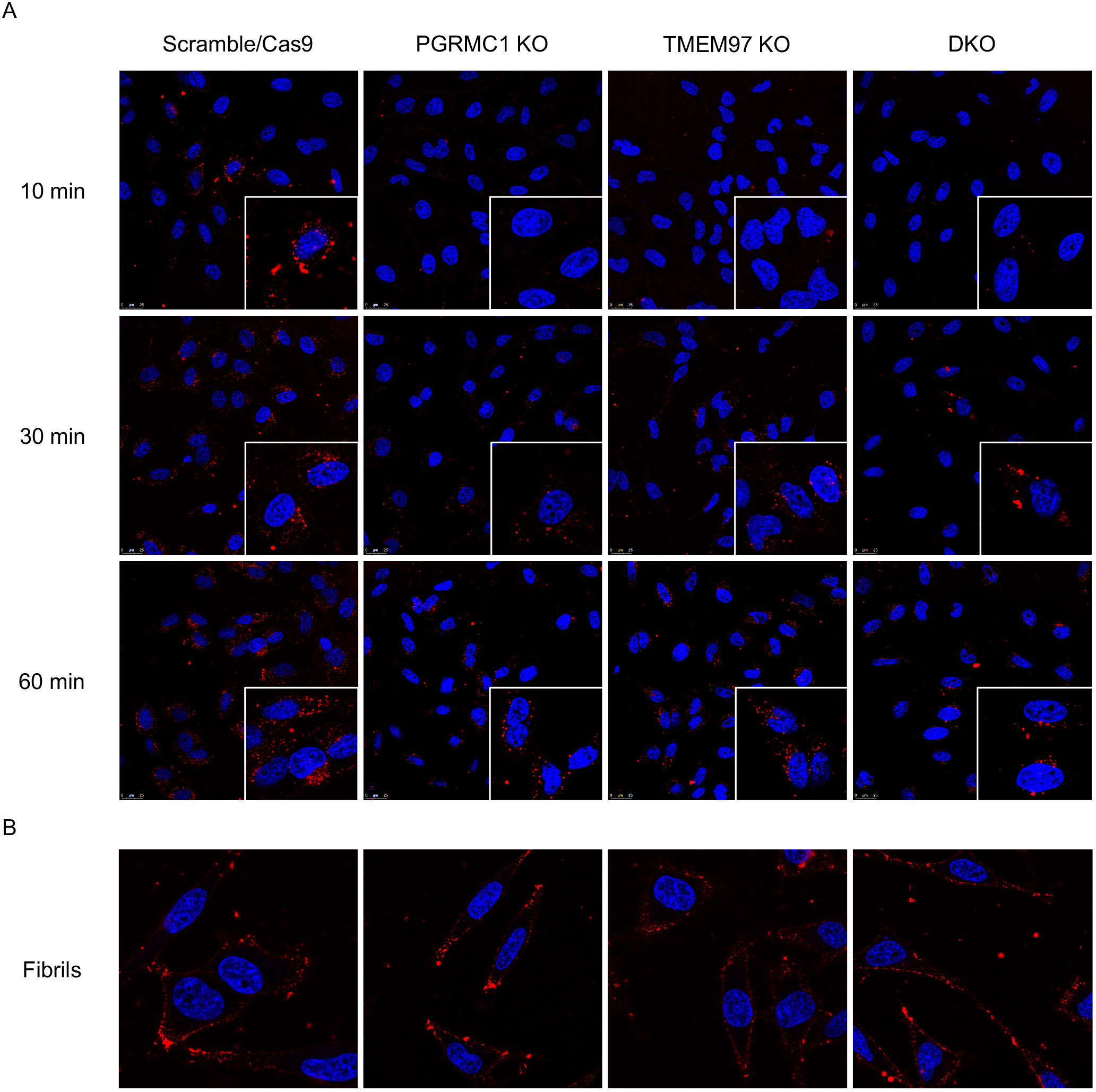
Confocal images of HeLa cell lines treated with fluorescent Aβ42 oligomers and fibrils. (A) Fluorescent Aβ42 oligomers treated at 10 min, 30 min, and 60 min. Inserts are expanded cells of interest. (B) Fluorescent Aβ42 treated for 60 minutes. Cells were stained with Hoechst (blue) prior to treatment with fluorescent Aβ42 (red) for indicated times, washed, and imaged immediately.

### TMEM97 and PGRMC1 primary rat cortical neurons

The next goal of this study was to assess the ability of TMEM97 and PGRMC1 ligands to inhibit the uptake Aβ42 in its various aggregation states and the apoE isoforms, apoE2, apoE3, and apoE4. Our first goal was to identify the presence of the trimeric complex in cultured primary neurons. For these studies we used mature DIV 21 primary rat cortical neurons, evident by their morphology and expression of MAP2 (Fig 3A). We identified the presence of the trimeric complex in a pairwise proximity ligation assay (Fig 3 C), as evident by the interaction of LDLR and PGRMC1, LDLR and TMEM97, and TMEM97 and PGRMC1. Taken together, these data indicate that the three proteins form a trimeric complex in primary neurons. Radiolabeled RHM-4 was able to bind with homogenates of primary rat neurons (Fig. 3 B), with an observed Kd of 1.37 nM and a Bmax of 1448 fmol/mg, further indicating the presence and high expression of TMEM97 in the primary neurons.

**Figure 3.**
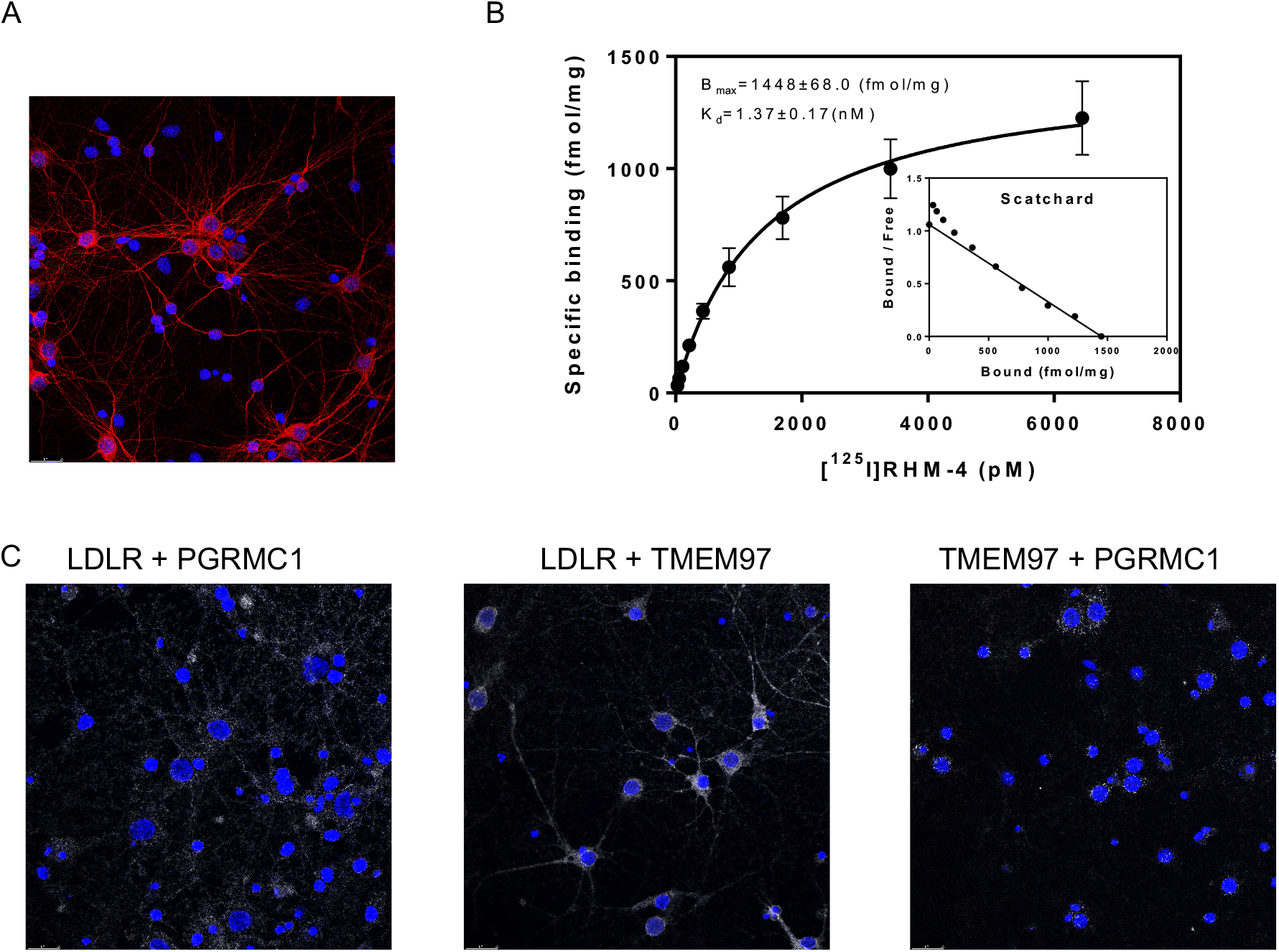
Characterization of Primary Neurons. (A) Confocal microscopy images of primary neuron MAP2 staining (red) indicates primary neurons are mature and display healthy neuronal structure. (B) Determination of sigma-2 receptor densities in rat cortical neurons by receptor binding assay using [^125^I]RHM-4. Inset is a scatchard plot. K_d_ and B_max_ were calculated by a nonlinear regression analysis using GraphPad Prism software. Data represent mean ± SD of at least 3 experiments performed in duplicate. (C) Pairwise Proximity Ligation Assay on DIV 21 primary rat cortical neurons. Confocal microscopy images of fluorescent signals (white) indicate an interaction between LDLR and PGRMC1, LDLR and TMEM97, and TMEM97 and PGRMC1; indicating the presence of a trimeric complex.

Primary neurons were treated with mAβ42 in the presence or absence of apoE2, apoE3, and apoE4, for 24 hours. For each treatment, a no-compound-treated group was compared to cells treated with 500nM RHM-4, SW43, and AG-205. The amount of Aβ42 and apoE was quantified via ELISA (Fig. 4 A and D). Results indicate that pharmacological inhibition of TMEM97 and PGRMC1 results in a decrease in mAβ42 and all apoE isoforms. There was an increase in uptake of Aβ42 in complex with the various apoE isoforms in a rank order of apoE2 < apoE3 < apoE4; interestingly all compounds were able to inhibit the mAβ42-ApoE complex with all three isoforms (E2, E4, E4), indicating potential therapeutic use in patients with any of the apoE alleles. The same trend was observed for oAβ42 in the presence and absence of the apoE isoforms (Fig 4 B and E). As expected from the previous observation that fAβ42 is not internalized but rather accumulates on the cell surface, fAβ42 was not readily taken up by cultured neurons, and the compounds did not have an effect on cell associated fAβ42 alone or in a complex with apoE. Curiously, apoE was not inhibited by the TMEM97 or PGRMC1 ligands when in a complex with fAβ42, which may be due to the fact that it accumulates on the surface along with fAβ42 and is not internalized. We observed that the internalized oAβ42 was associated with MAP2 positive neurons (SI Appendix, Fig. S2).

**Figure 4.**
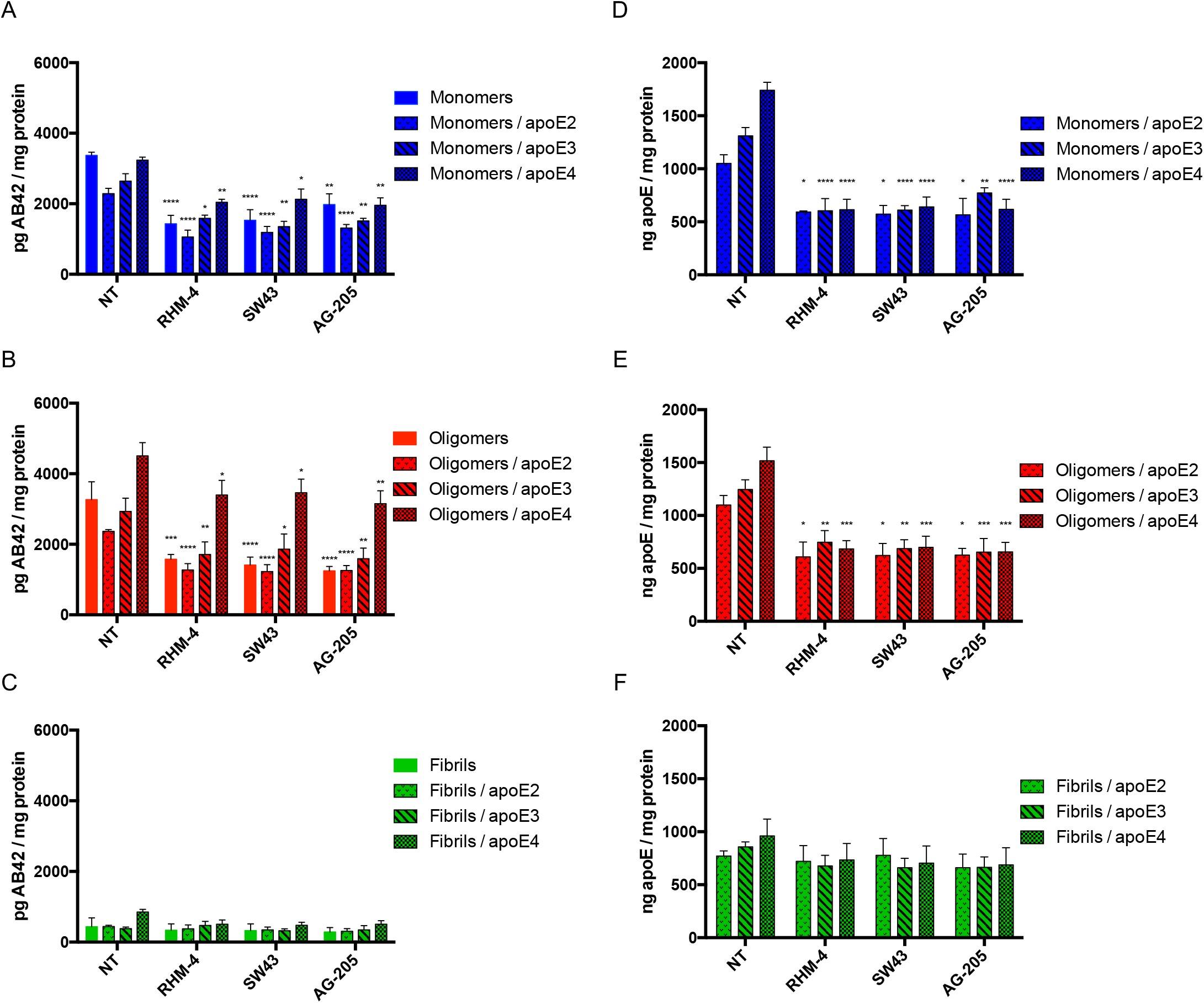
Treatment of primary rat cortical neurons (DIV21) with various sigma 2 ligands reduces the uptake of Aβ42 monomers and oligomers in the presence and absence of apoE. Primary neurons were treated with 500nM RHM-4, SW43, or AG-205 and uptake of Aβ42 monomers, oligomers, and fibrils with or without apoE2, apoE3 or apoE4 was quantified. Uptake was compared to no compound treated controls (NT). Drug treatment resulted in a reduction of Aβ42 (A) monomers and (B) oligomers but not (C) fibrils when treated alone or in a complex with any apoE isoforms. (D-F) Quantification of apoE was assessed for these treatment groups. Cell associated Aβ42 and apoE was determined via ELISA of cell lysates normalized to total cell protein. Data represents mean ± SEM (n=4); *, p < 0.05; **, p < 0.01; ***, p < 0.001; **** p < 0.0001; one way ANOVA for each treatment compared to control.

The next step was to determine if the TMEM97/PGRMC1/LDLR trimeric complex is present in human brain tissue. In this study, samples of frontal cortex from postmortem cognitively normal subjects and a subject identified with AD (Fig. 5) were used. The normal brain tissue section was from a patient with no clinical history of dementia or cognitive impairment, and the presence of neurofibrillary tangles was not observed in the pathology report. The AD brain section was from a patient with a clinical history of AD, with the pathology report indicating high level of amyloid beta, tau, and alpha-synuclein positive inclusions, along with numerous neurofibrillary tangles prominent in the frontal cortex. The AD tissue section displayed positive staining for Aβ whereas the normal human tissue sections did not show a signal in immunofluorescent staining with an Aβ antibody (SI Appendix, Fig. S3). We observed that the trimeric complex is intact in both human tissue samples, and signifies that our results in both the HeLa model system and primary cultured neuronsative of the conditions present in adult human brain. Consequently, our observations that pharmacologically targeting this complex blocks the uptake of Aβ42 and ApoE-associated Ab42 may represent a novel strategy for preventing the cellular uptake and cell-to-cell transmission of beta amyloid peptide.

**Figure 5.**
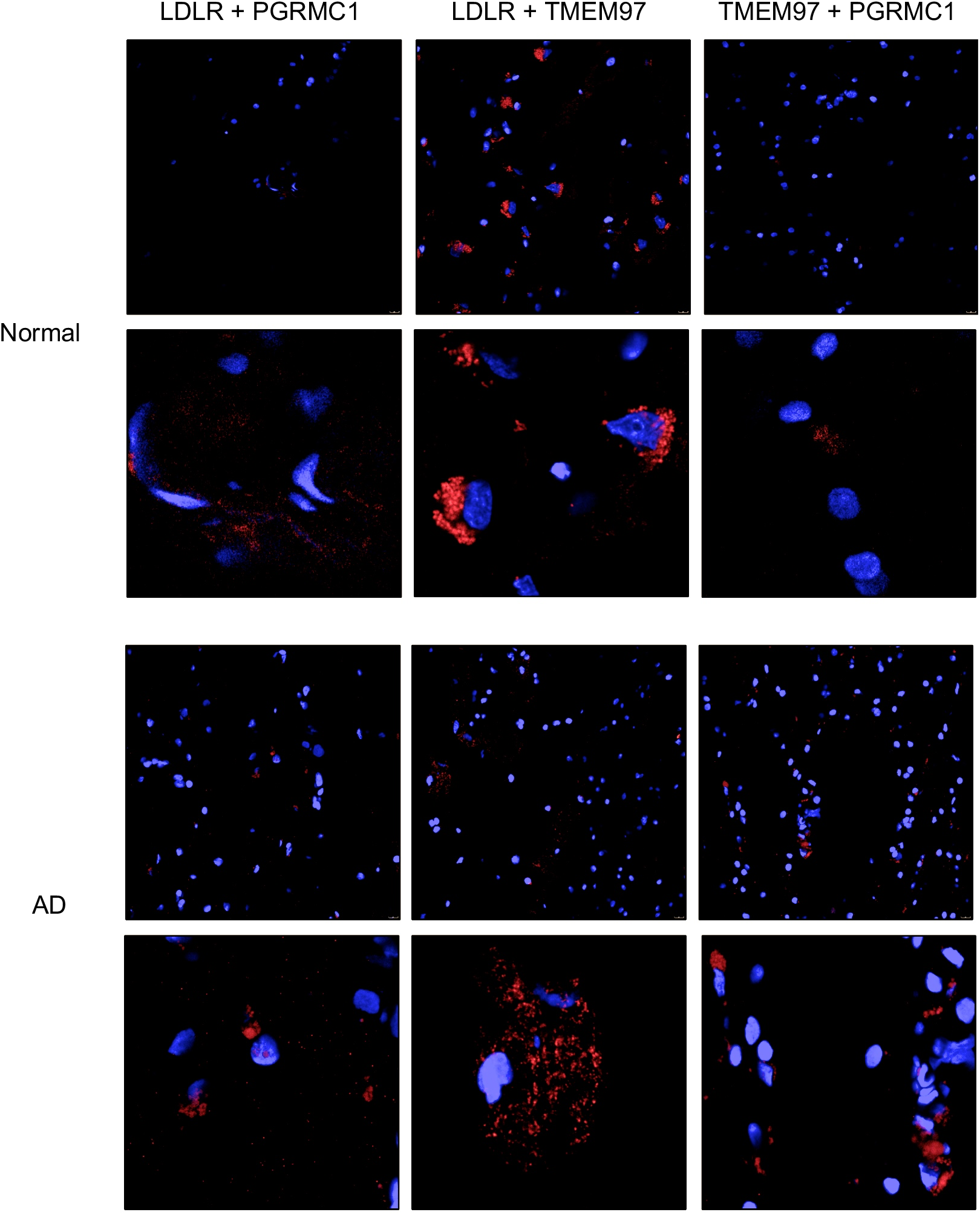
Pairwise Proximity Ligation Assay on normal adult human brain and Alzheimer’s Disease human brain. Fluorescent signals (red) indicate an interaction between LDLR and PGRMC1, LDLR and TMEM97, and TMEM97 and PGRMC1; indicating the trimeric complex is intact in normal and diseased human brains. Top 40X magnification, Bottom, expanded field of interest.

## DISUCSSION

In the current study, we provide strong experimental evidence that TMEM97 and PGRMC1 are therapeutic targets for inhibition of Aβ42 and apoE uptake. Evidence suggests that increased uptake of these neurotoxic Aβ42 peptides results in accumulation and aggregation within neurons eventually leading the formation of plaques and neuronal death [34, 35]. Due to the apoE4 allele being greatest risk factor associated with development of AD, LDLR and the LRP family have been the focus of many studies [36–38]. In our previous study, we described a trimeric protein complex formed between LDLR, TMEM97, and PGRMC1, and that this intact complex is important for efficient uptake of LDL in cancer cells. In the current study, we confirmed that the trimeric TMEM97/PGRMC1/LDLR complex is intact in primary rat cortical neurons (Fig. 3C) and in human brain tissues, both in non-diseased human brain and in AD brain samples (Fig. 5). The goal of this study was to assess the importance of this intact complex, and its potential target for pharmacological intervention, on Aβ42 and apoE-associated Aβ42 uptake.

Knocking out TMEM97 and/or PGRMC1 in HeLa cells resulted in a marked decrease of mAβ42 and oAβ42 uptake alone or when in a complex with apoE. The uptake of mAβ42 in HeLa cells was enhanced by the presence of apoE (Fig. 1 A and C), but interestingly this reliance on apoE as a chaperone for uptake was not observed in the primary cultured neurons (Fig. 4 A and D). Conversely, oAβ42 did not require apoE to be taken up by HeLa cells (Fig. 1 A and C) or primary neurons (Fig. 4 B and E). These observations are consistent with previous reports [39]. In both the HeLa model system and primary neuronal cells, RHM-4, SW43, and AG-205 were able to reduce uptake of mAβ42 and oAβ42 alone and when in a complex with apoE. The results highlight the role of TMEM97 and PGRMC1 in neuronal Aβ42 and apoE-associated Aβ42 uptake, and reinforces the therapeutic potential of the TMEM97-PGRMC1-LDLR trimeric complex as a target for reducing Aβ42 accumulation in neurons.

The propagation of toxic aggregates throughout the AD brain has spurred research on the neuronal transfer of Aβ42. This cell-to-cell transfer was initially explored after the observation that the neuronal dysfunction that led to memory impairment also caused pathological changes in neighboring cells [25, 40]. There have been several proposed mechanisms by which this process occurs including exosomes and direct transmission of Aβ42 from neuron to neuron that are directly connected by tunneling nanotubes [22]. Exosomes have been investigated as a means of neuron to neuron transfer of Aβ42, as oAβ42 has been detected within neuronal exosomes. Data from cells treated with these exosomes also revealed that the exosomes facilitated delivery of oAβ42 and induced toxic effects [24]. It is tempting to speculate that LDLR or LRP1 play a role in the uptake of these exosomes, thus implicating the TMEM97-PGRMC1-LDLR complex as a potential target in reducing the spread of cytotoxic aggregates. Further research delineating the role of TMEM97 and PGRMC1 in neuronal transfer of not only aggregated Aβ42 species, but other aggregated protein such as alpha synuclein and tau, will be of great interest.

In conclusion, the results from this study reveal that TMEM97 and PGRMC1 may be novel therapeutic target for inhibiting neuronal uptake of Aβ42 and ApoE-associated Aβ42 and potentially reduce Aβ-associated neurotoxicity. The modulation of intercellular and extracellular Aβ42 pools may have a positive influence on clearance by keeping Aβ42 in the extracellular space where it may be readily available for proteolytic degradation or clearance to the CSF. Developing compounds that target the TMEM97/PGRMC1/LDLR complex may represent a novel strategy in delaying the clinical progression of AD. Whether or not this mechanism has implications in other CNS disorders involving protein aggregates such as alpha-synuclein in Parkinson’s Disease or the aggregation of tau in tauopathies is currently under investigation.

## MATERIALS AND METHODS

### Materials

*N*-(4-(6,7-dimethoxy-3,4-dihydroisoquinolin-2(1H)-yl)butyl)-2,3-dimethoxy-5-iodo-benzamide and [125I]*N*-(4-(6,7-dimethoxy-3,4-dihydroisoquinolin-2(1H)-yl)butyl)-2,3-dimethoxy-5-iodo-benzamide were synthesized as previously described [41, 42]. Hoechst 33342 (BD Pharmingen, 561908).

### Postmortem Human Brain Samples

Human brain tissue samples from donor subjects following neuropathological evaluation were obtained from the brain bank at the Center for Neurodegenerative Disease Research at the University of Pennsylvania based on previously published criteria [43–45]. The methodology for brain harvest, selection of areas of interest, and diagnostic procedures and criteria have been previously reported.

### Cell Culture

HeLa cell TMEM97 knockout, PGRMC1 knockout, and DKO cell lines were generated as previously described [8]. Cells were cultured in MEM with 10% FBS, 1X penicillin/streptomycin, 2mM L-glutamine, and 1X MEM non-essential amino acids. For uptake experiments, cells were plated, and incubated for 24 hours, media was removed and cells were incubated in MEM containing 10% lipoprotein depleted serum for an additional 24 hours prior to treatment.

### Primary Cortical Rat Neuron Cell Culture

Primary neurons were acquired from University of Pennsylvania Neurons R Us and plated in poly-D-lysine coated 6 well plates at 400 cells/well or in poly-D-lysine coated 8 well chamber slides at 100 cells/well in Neurobasal Plus Media supplemented with B27 Plus, 1X GlutaMAX, and 0.5X penicillin/streptomycin. Media was partially exchanged every three days. Cells were cultured for 21 days prior to use in experiments.

### Aβ42 fibril preparation

Aβ42 fibril was prepared as described previously [32]. Briefly, monomeric Aβ42 (Millipore-Sigma, A9810) was dissolved in hexafluoroisopropanol (HFIP) at a concentration of 2 mg/mL and incubated for 1 hour at 37 °C until the peptide completely dissolved. Then HFIP was evaporated under air. The peptide powder was dissolved again in HFIP (2 mg/mL), aliquoted (50 μL) and left to dry overnight under vacuum. Aliquots were stored in a freezer at −20 °C.

To prepare fibrils, the HFIP-treated peptide aliquot was dissolved in 10 mM NaOH (10 μL) solution, then the sample was diluted with 90 μL 10 mM phosphate buffer, pH 7.4. Concentration of the peptide solution was confirmed by measuring the absorbance at 214 nm with NanoDrop 2000c spectrophotometer. Extinction coefficient (76848 M^−1^cm^−1^) was calculated using literature values [30]. Next, the solution was agitated by a continuous slow rotation at room temperature for 3 days and fibril formation was confirmed by TEM.

Oligomers were prepared as previously described [31]. An aliquot of HFIP-treated peptide was dissolved in anhydrous DMSO (5 μM) and diluted to 100 μM in serum-free and phenol red-free cell culture media, followed by incubation at 4 °C for 24 hours.

Fluorescent Aβ42 oligomers and fibrils were prepared similarly with HiLyte Fluor 555-labeled Aβ42 (Anaspec, AS-60480-01) and mixed with non-labeled Aβ42 peptides in a 1:2 ratio.

### Transmission electron microscopy

Samples (5 μL) were spotted onto glow-discharged formvar/carbon-coated, 200-mesh copper grids (Ted Pella). After 1 min, grids were washed briefly with water and stained with one 10 μL drop of 2% w/v uranyl acetate for 1 min. The excess stain was removed by filter paper and the grids were dried under air. Samples were imaged with a Tecnai FEI T12 electron microscope at an acceleration voltage of 120 kV. Images were recorded on a Gatan OneView 4K Cmos camera.

### ApoE Preparation

Recombinant apoE2 (Millipore-Sigma, SRP4760), apoE3 (Millipore-Sigma, SRP4696), and apoE4 (Abcam, ab50243) was reconstituted per manufacture instruction to a concentration of 1 mg/mL, aliquoted, and stored at −80 °C. Prior to use, apolipoprotein was lipidated with dimyristoyl- phosphatidylcholine (DMPC) (Millipore-Sigma, P2663) using a procedure adapted from [46]. Briefly, DMPC vesicles (10 mg/mL in in PBS) were prepared by sonicating until solution was clear. These were mixed with apoE (37.5 μL vesicles to 100 μg apoE) and the solution was cycled three times through the transition temperature of the DMPC (23.5 °C) by warming to 30 °C for 30 minutes and cooling to 15 °C for 30 minutes using a thermocycler. To form a complex with Aβ42, the lipidated apoE solution was added immediately to Aβ42 (at a molar ratio of 1:4) and incubated at 37 °C for 30 minutes.

### Dot blot analysis

2 μg of Aβ42 monomers, oligomers, and fibrils were spotted on nitrocellulose membrane and allowed to dry for 1 hour at room temperature. Membranes were blocked with Odyssey blocking buffer (927–40100 Licor Biotechnology, Lincoln, NE) for 1 hour at room temperature, then incubated with primary antibody (A11 1:10000) (Invitrogen, AHB0052) or 6E10 (1:1000) (Biolegend, 803001) overnight at 4°C. Membranes were washed 3 times with PBS-T (0.2% Tween-20) for 5 minutes, then incubated with secondary antibody secondary antibody, IRDye 680CW anti-rabbit IgG (Licor Biotechnology, 926–68071) or IRDye 800CW anti-Mouse IgG (Licor Biotechnology, 926-32210) at a 1:8,000 dilution for 1 hour at room temperature. Membranes were washed 3 times with PBS-T for 5 minutes, followed by a rinse in PBS, then imaged using Odyssey CLx Infrared Imaging System (Licor Biotechnology, Lincoln, NE).

### Aβ42 and apoE ELISA

To quantify Aβ42 and apoE uptake, cells were treated, washed with PBS 3 times, then lysed in RIPA buffer (sc-24948 Sigma-Aldrich, St. Louis, MO, USA) supplemented with protease inhibitor cocktail, and phosphatase inhibitor cocktail 2 and 3 (sc-24948 Sigma-Aldrich, St. Louis, MO, USA). Total Aβ42 was measured via ELISA (Thermo Fisher, KHB3544), total apoE was quantified via ELISA (Abcam ab108813). Uptake was normalized to total mg of protein per sample measured via Bio-Rad DC protein assay kit (5000112 Bio-Rad Laboratories, Hercules, CA, USA).

### Confocal Microscopy

Cells were plated in 8-well chamber slides (Lab-Tek cc2 plates, 154534PK Thermo Scientific). Cells were washed 3 times with PBS then fixed with 4% paraformaldehyde (Santa Cruz) for 10 minutes at room temperature, washed 3 times with PBS, then permeabilized with 0.1% Triton X-100 in PBS for 10 minutes at room temperature. Cells were blocked with 10% Goat Serum (50062Z Thermo Scientific) for one hour then incubated with goat anti-PGRMC1 (Abcam ab48012) 1:200 in PBST +1% goat serum overnight, washed 3 times with PBST, then incubated with 1:200 donkey anti-goat Alexa488 secondary antibody (A-11055 Invitrogen) in PBST for 1 hour. Cells were then washed 3 times in PBST and blocked with 10% goat serum for an hour, then incubated with 1:50 mouse anti-LDLR (Novus NBP1-78159) in PBST +1% goat serum overnight, washed 3 times with PBST, then incubated with 1:200 goat anti-mouse Cy3 secondary antibody (A10521 Invitrogen) in PBST for 1 hour. Cells were washed 3 times with PBST and blocked with 10% goat serum for an hour, then incubated with rabbit anti-TMEM97 primary antibody (Novus NBP1-30436) 1:200 in PBST +1% BSA overnight, washed 3 times with PBST, then incubated with goat anti-rabbit Alexa568 secondary antibody (A-11011 Invitrogen) 1:200 in PBST for an hour. Cells were washed 3 times in PBST, once in PBS, then mounted in ProLong Glass antifade mounting media (Thermo Fisher, P36980). Images were acquired at 40X magnification on a Leica STED 8X Super-resolution Confocal Microscope.

### Proximity Ligation Assays

For the Proximity Ligation Assay, the Duolink In Situ PLA Far Red kit was followed according to manufacturer instructions (Sigma DUO92105).

### Sigma-2 receptor saturation binding assay

For culturing primary cortical neurons, cortical cell suspensions were plated onto poly L-lysine (PLL)-coated T-75 flasks at 4×10^6^ cells per flask and cultured in neurobasal medium supplemented with B27 (1x), 100 units/ml penicillin, and 100 μg/ml streptomycin. The cell culture medium was partially replaced every 7 days. 2 week-old neurons were harvested by scraping and centrifugation at 2500×g for 20 min at 4°C. Cell pellets were frozen at −80 °C until use.

For cell membrane preparation, the cell pellets were re-suspended in 10 mL ice-cold Phosphate-Buffered Saline (PBS), and homogenized using Wheaton overhead stirrer (120 Vac Overhead Stirrer, Millville, NJ) at the speed of 2 for 30 sec. Cell homogenates were then centrifuged at 31,000 × g for 20 min at 4°C. The pellets were re-suspended in 1 mL ice-cold PBS and stored at −80°C freezer.

For saturation binding assays, cell membranes were incubated with [^125^I]RHM-4 (0.02 - 9 nM) for 90 min at room temperature. After incubation, the bound ligands were filtrated and collected on glass fiber papers (Whatman grade 934-AH, GE Healthcare, Pittsburgh, PA) with a M-24 Brandel filtration system (Brandel, Gaithersburg, MD), and counted with a Wizard2 Automatic Gamma Counter 2470. Nonspecific binding was determined in the presence of 10 μM DTG. The K_d_ and B_max_ values were calculated by a nonlinear regression analysis using GraphPad Prism version 6 (GraphPad, La Jolla, CA). Protein concentrations were determined by Lowry method.

## LIST OF ABBREVIATIONS

Aβ42: amyloid beta 1-42
AD: Alzheimer’s Disease
APP: amyloid precursor protein
apoE: apolipoprotein E
CNS: central nervous system
DKO: double knockout
DMPC: dimyristoyl-phosphatidylcholine
fAβ42: Aβ42 fibrils
HFIP: hexafluoroisopropanol
LDL: low density lipoprotein
LDLR: Low Density Lipoprotein Receptor
LPDS: lipoprotein depleted serum
LRP: LDLR-related protein
mAβ42: Aβ42 monomers
oAβ42: Aβ42 oligomers
PBS: phosphate buffered saline
PBS-T: phosphate buffered saline with 0.2% Tween 20
PGRMC1: Progesterone Receptor Membrane Component 1
PLL: poly L-lysine
TEM: transmission electron microscopy
TMEM97: Transmembrane Protein

## DECLARATIONS

### Ethics approval and consent to participate

All human tissue was used in accordance with the University of Pennsylvania IRB protocol.

### Consent for publication

Not Applicable

### Availability of data and materials

Not Applicable

### Authors’ contributions

AR performed and analyzed uptake experiments in HeLa cells and primary rat neurons, ELISA analysis, confocal microscopy experiments, and manuscript preparation. ZL performed and analyzed TEM experiments and Aβ42 fibril preparations, CZ performed CRISPR knockout of HeLa cell lines and Aβ42 microscopy on human tissues, CW performed and analyzed radioligand binding experiments, VL and JT were responsible for tissue sample acquisition, analysis, and pathology, RHM was responsible for experimental organization and rationale, data analysis, organization, and manuscript preparation. All authors read and approved the final manuscript.

## Acknowledgements

Not applicable

