## Supplementary Figures and Legends for "The Sigma-2 Receptor/TMEM97, PGRMC1, and LDL Receptor complex are responsible for the cellular uptake of Aβ42 and its protein aggregates"

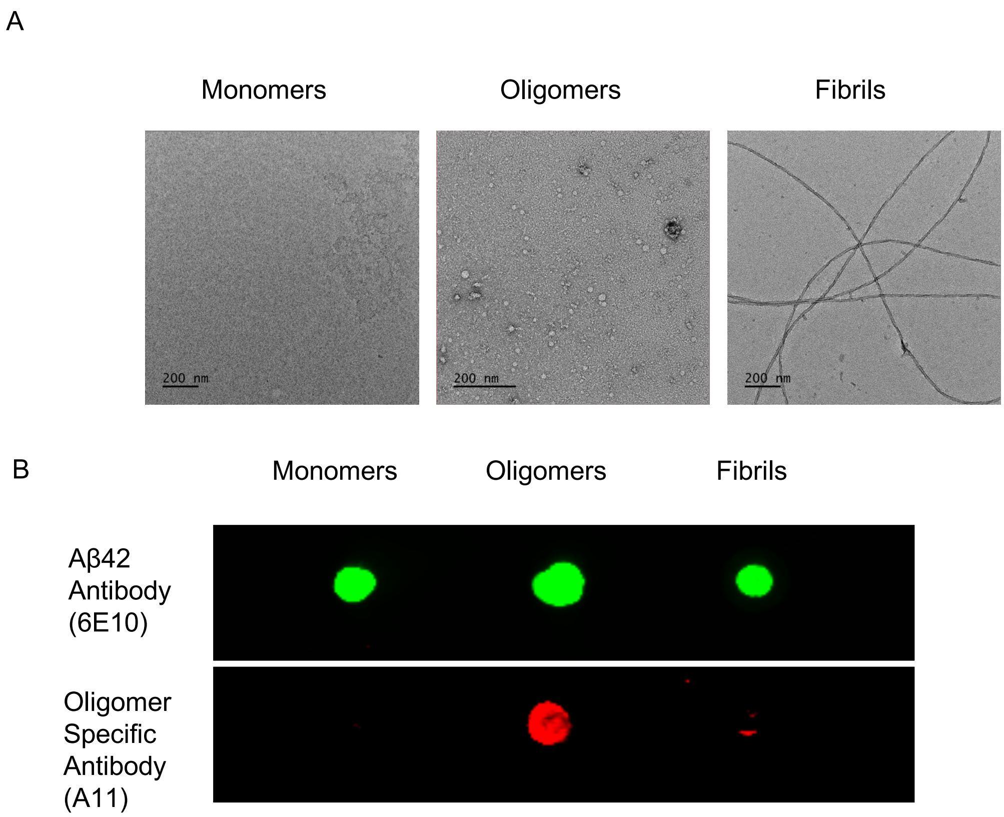


Supplementary Figure S1. Characterization of Aβ42 samples. (A) Negative-stained TEM images of Aβ42 monomers, oligomers, and fibrils. (B) Dot blot analysis indicates mAβ42, oAβ42, and fAβ42 all show positive reacitivity for Aβ42 6E10 antibody, but only oAβ42 showed positive reactivity to the oligomer specific antibody A11.

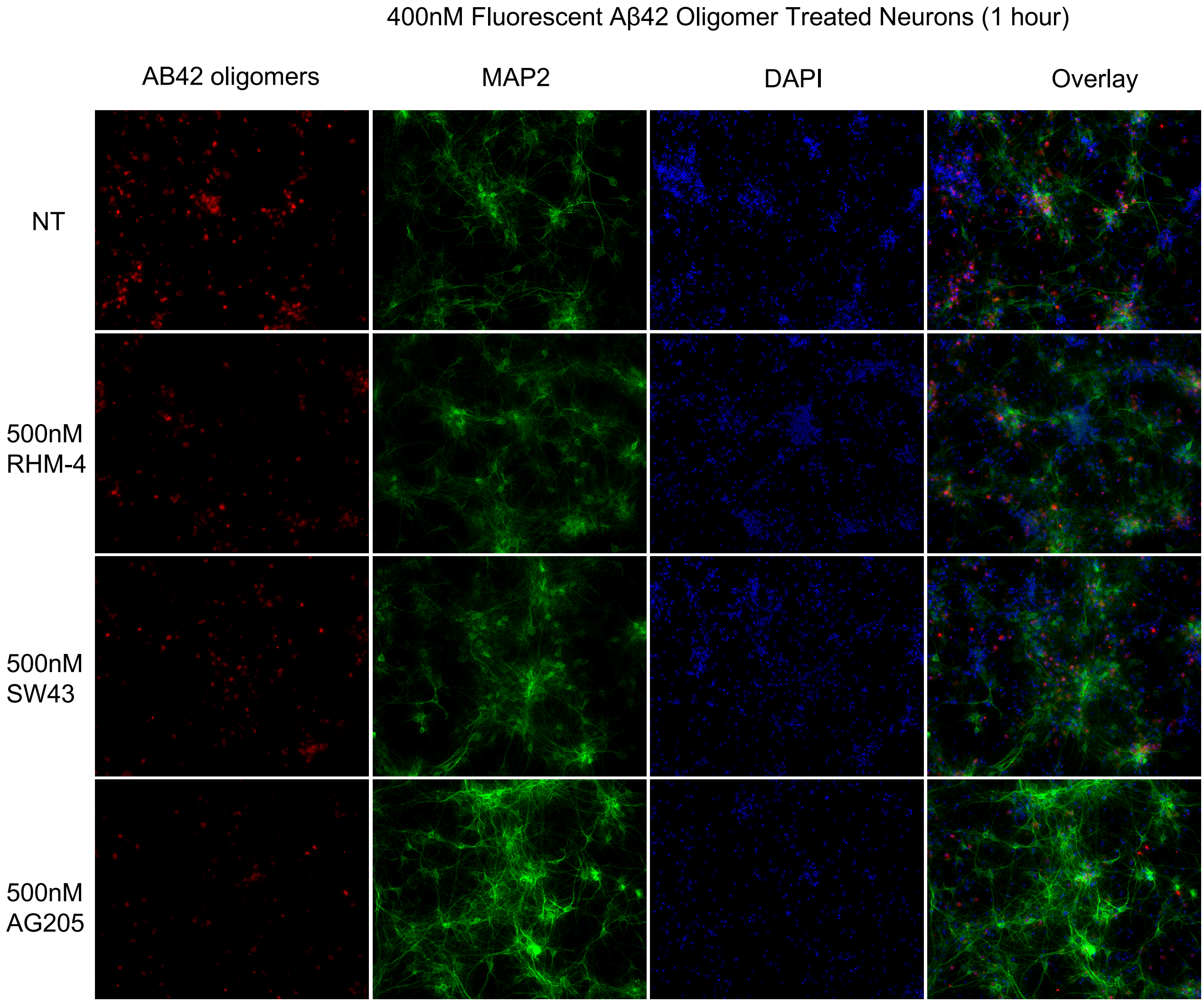


Supplementary Figure S2. Uptake of fluorescently labeled Aβ42 oligomers is by primary rat cortical neurons (DIV21) is reduced in the presence of sigma 2 ligands. Primary neurons were treated with 400 nM Aβ42 oligomers for 2 hours at 37 °C, fixed and stained with MAP2. The control compound treatment group (NT) showed higher uptake than the compound treated groups. Aβ42 signal (red) was associated with MAP2 positive neurons (green).

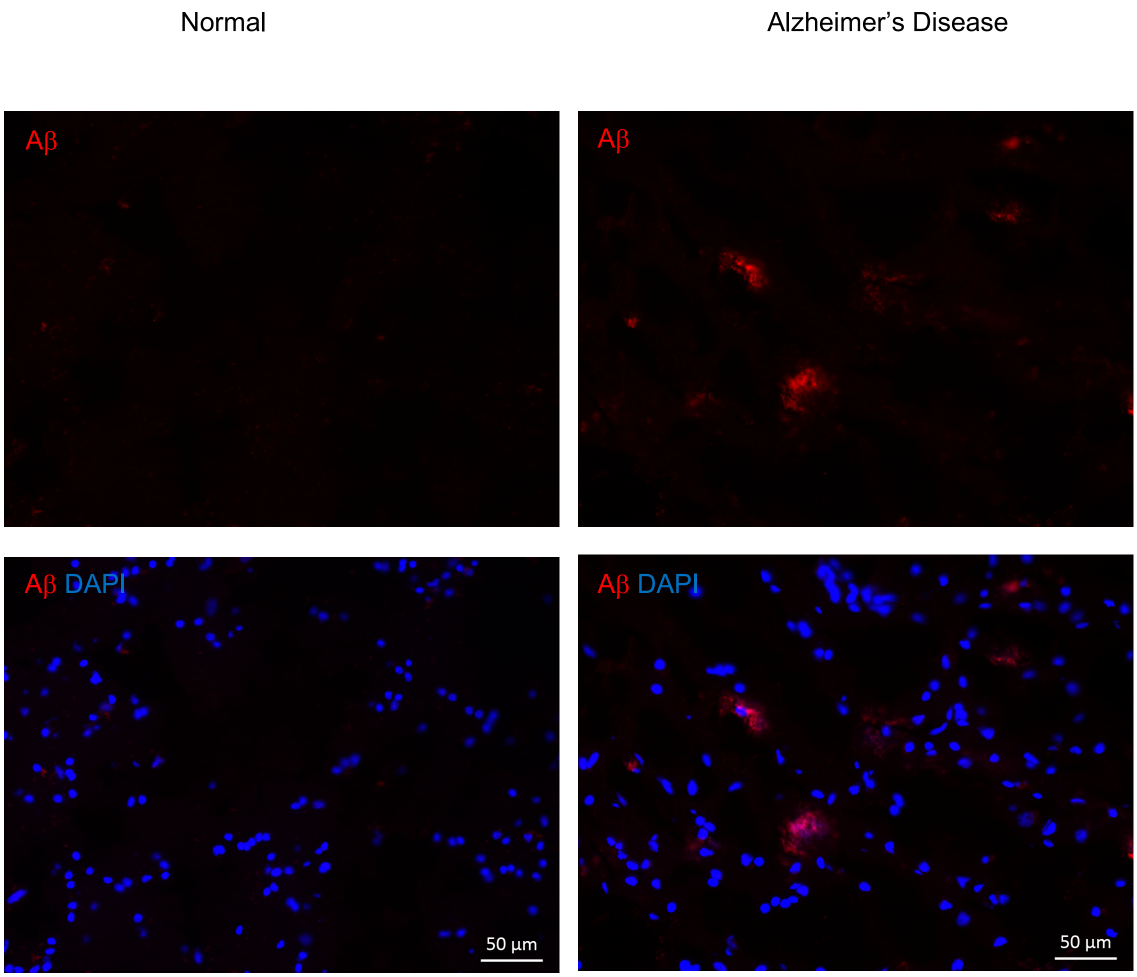


Fig. S3 Immunofluorescence staining of Aβ in normal and AD human brain cortex. Aβ staining shows positive signal in Alzheimer’s Disease human brain tissues but not in normal human brain tissues.
